# Coherent olfactory bulb gamma oscillations arise from coupling independent columnar oscillators

**DOI:** 10.1101/213827

**Authors:** Shane T. Peace, Benjamin C. Johnson, Jesse C. Werth, Guoshi Li, Martin E. Kaiser, Izumi Fukunaga, Andreas T. Schaefer, Alyosha C. Molnar, Thomas A. Cleland

## Abstract

Spike timing-based representations of sensory information depend on embedded dynamical frameworks within neuronal networks that establish the rules of local computation and interareal communication. Here, we investigated the dynamical properties of olfactory bulb circuitry in mice of both sexes using microelectrode array recordings from slice and *in vivo* preparations. Neurochemical activation or optogenetic stimulation of sensory afferents evoked persistent gamma oscillations in the local field potential. These oscillations arose from slower, GABA(A) receptor-independent intracolumnar oscillators coupled by GABA(A)-ergic synapses into a faster, broadly coherent network oscillation. Consistent with the theoretical properties of coupled-oscillator networks, the spatial extent of zero-phase coherence was bounded in slices by the reduced density of lateral interactions. The intact *in vivo* network, however, exhibited long-range lateral interactions that suffice in simulation to enable zero-phase gamma coherence across the olfactory bulb. The timing of action potentials in a subset of principal neurons was phase-constrained with respect to evoked gamma oscillations. Coupled-oscillator dynamics in olfactory bulb thereby enable a common clock, robust to biological heterogeneities, that is capable of supporting gamma-band spike synchronization and phase coding across the ensemble of activated principal neurons.

**New & Noteworthy:** Odor stimulation evokes rhythmic gamma oscillations in the field potential of the olfactory bulb, but the dynamical mechanisms governing these oscillations have remained unclear. Establishing these mechanisms is important, as they determine the biophysical capacities of the bulbar circuit to, for example, maintain zero-phase coherence across a spatially extended network, or coordinate the timing of action potentials in principal neurons. These properties in turn constrain and suggest hypotheses of sensory coding.

## Introduction

Gamma-band field potential oscillations reflect action potential timing constraints in assemblies of principal neurons. As such, they constitute side effects of timing-based neural codes, which are able to communicate statistically reliable information much more rapidly than rate-based codes and have clearer postsynaptic mechanisms of operation (Fries 2009, Shouval et al 2010, Theunissen & Miller 1995). Gamma-band rhythmicity is generated within the earliest central circuits of the olfactory system, imposing temporal structure on rate-coded primary sensory inputs and regulating action potential timing in second-order principal neurons. Disruption of these rhythms impairs olfactory performance (Kashiwadani et al 1999, Lepousez & Lledo 2013, Stopfer et al 1997). In the mammalian olfactory bulb (OB), gamma oscillations are intrinsic, arising from synaptic interactions between glutamatergic mitral and projecting tufted cells (MCs; principal neurons) reciprocally connected with GABAergic granule cells (GCs) and potentially other interneurons within the OB external plexiform layer (EPL) (Cleland 2014, Fukunaga et al 2014, Huang et al 2013, Lagier et al 2004, Lepousez & Lledo 2013, Mori et al 2013). These oscillations persist *in vivo* when cortical connections are severed (Martin et al 2004, Martin et al 2006, Neville & Haberly 2003), and have been transiently evoked in OB slices *in vitro* (Friedman & Strowbridge 2003, Gire & Schoppa 2008, Lagier et al 2004, Schoppa 2006). These data underlie current hypotheses that OB circuitry, in addition to processing the *information content* of primary sensory inputs, also transforms the *metric* of this sensory information from a rate-coded basis among primary olfactory sensory neurons (OSNs) into a sparser, spike timing-based representation at the output of the OB (Cleland & Borthakur 2020, Linster & Cleland 2010, Litaudon et al 2008, Luna & Schoppa 2008, McIntyre & Cleland 2016, Poo & Isaacson 2009), thereby gaining the attendant advantages of this coding scheme (Fries 2009).

Implicit in these theories of timing-based computation in the OB is the principle that gamma oscillations are substantially coherent across the entire olfactory bulb (Freeman 1978, Kay & Lazzara 2010, Linster & Cleland 2010). More broadly, the theory of communication through coherence presumes zero-phase gamma coherence within intra-areal populations (Bastos et al 2015, Fries 2015). However, the capacity for such broad coherence in OB is severely challenged by the substantial heterogeneity in mitral cell activation levels that encodes odor identity, and by other limitations of the dynamical systems thought to underlie OB oscillations. By what mechanisms, then, is the EPL network able to both preserve essential coding heterogeneity across principal neurons while also maintaining broadly coherent activity across the entire OB?

To elucidate these mechanisms, we developed an OB slice preparation that exhibits persistent gamma oscillations in response to either the optogenetic excitation of OSN arbors or transient pharmacological stimulation with metabotropic glutamate receptor (mGluR) or cholinergic agonists, and recorded local field potentials (LFPs) across 59 discrete electrodes using a 60-electrode planar multielectrode array (MEA). Action potential timing in a subset of principal neurons was constrained with respect to the phase of these gamma oscillations, as has been previously observed (Bathellier et al 2006, Eeckman & Freeman 1990, Kashiwadani et al 1999). Notably, in contrast to existing findings utilizing a single large electrode to record LFPs (Bathellier et al 2006, Lagier et al 2004), the blockade of GABA(A) receptors did not reduce oscillatory power in the gamma band. Instead, our spatially distributed MEA electrodes revealed that GABA(A) blockade modestly reduced the peak oscillation frequency and decoupled neighboring areas of the OB, reducing the spatial extent of LFP coherence. Critically, this decoupling effect would appear as a reduction in gamma power when measured in single electrodes with broad pickup fields (see *Discussion*), which explains the difference between previous results and those described herein. We refer to the localized, decoupled regions of GABA(A) blockade-resistant oscillatory coherence as *columns.* It is likely that these columns each correspond to an individual glomerulus, the principal neurons innervating it, and the local interneurons affecting those principal neurons; accordingly, we suggest that MC resonance properties (Desmaisons et al 1999, Rubin & Cleland 2006), locally synchronized via intraglomerular gap junctions (Pouille et al 2017), underlie local, intracolumnar LFP oscillations. These separate columnar oscillators then are coupled with other columns via their mutual synaptic interactions with GABA(A)ergic granule cells within the EPL, thereby synchronizing MCs (Schoppa 2006), increasing the underlying oscillation frequency, and enabling intercolumnar coherence across the spatial extent of the EPL network (Li & Cleland 2017). Activation of the OB network (e.g., by odor stimulation) consequently is able to generate a selective and coherent assembly of activated principal neurons unconstrained by spatial proximity. To map these findings onto *in vivo* conditions, we then performed intracellular recordings paired with optogenetic stimulation both in slices and *in vivo* to determine the intercolumnar connectivity afforded by the extensive lateral dendritic projections of MCs, and employed a biophysical computational model of the OB to illustrate how the results from the slice experiments generalize to the intact OB *in vivo*.

## Methods

### Mouse lines

For MEA recordings, we used OMP-ChR2-EYFP (ORC-M; C57Bl/6- and 129-derived) plasmid transgenic mice of both sexes (Dhawale et al 2010), originally provided by Venkatesh Murthy, Harvard University. No sex differences were observed in recordings. For intracellular recordings, we used heterozygous Omptm1.1(COP4*/EYFP)Tboz/J knock-in mice of both sexes (Smear et al 2011), provided by Tom Bozza, Northwestern University. Each of these transgenic lines coexpresses channelrhodopsin-2 (ChR2) and enhanced yellow fluorescent protein (EYFP) in olfactory sensory neurons (OSN) under the control of the olfactory marker protein (OMP) promoter, hence limiting transgene expression within the OB to OSN axon terminals in the glomerular layer (Fig. 1*A*; also see (2013)). Additional MEA experiments were performed with CD-1 outbred mice (Charles River, Kingston, NY, USA) where noted. All procedures for MEA experiments were performed under the auspices of a protocol approved by the Cornell University Institutional Animal Care and Use Committee (IACUC). Cornell University is accredited by The Association for Assessment and Accreditation of Laboratory Animal Care (AAALAC International). All procedures for intracellular recording experiments were performed in compliance with German animal welfare guidelines.

**Figure 1.**
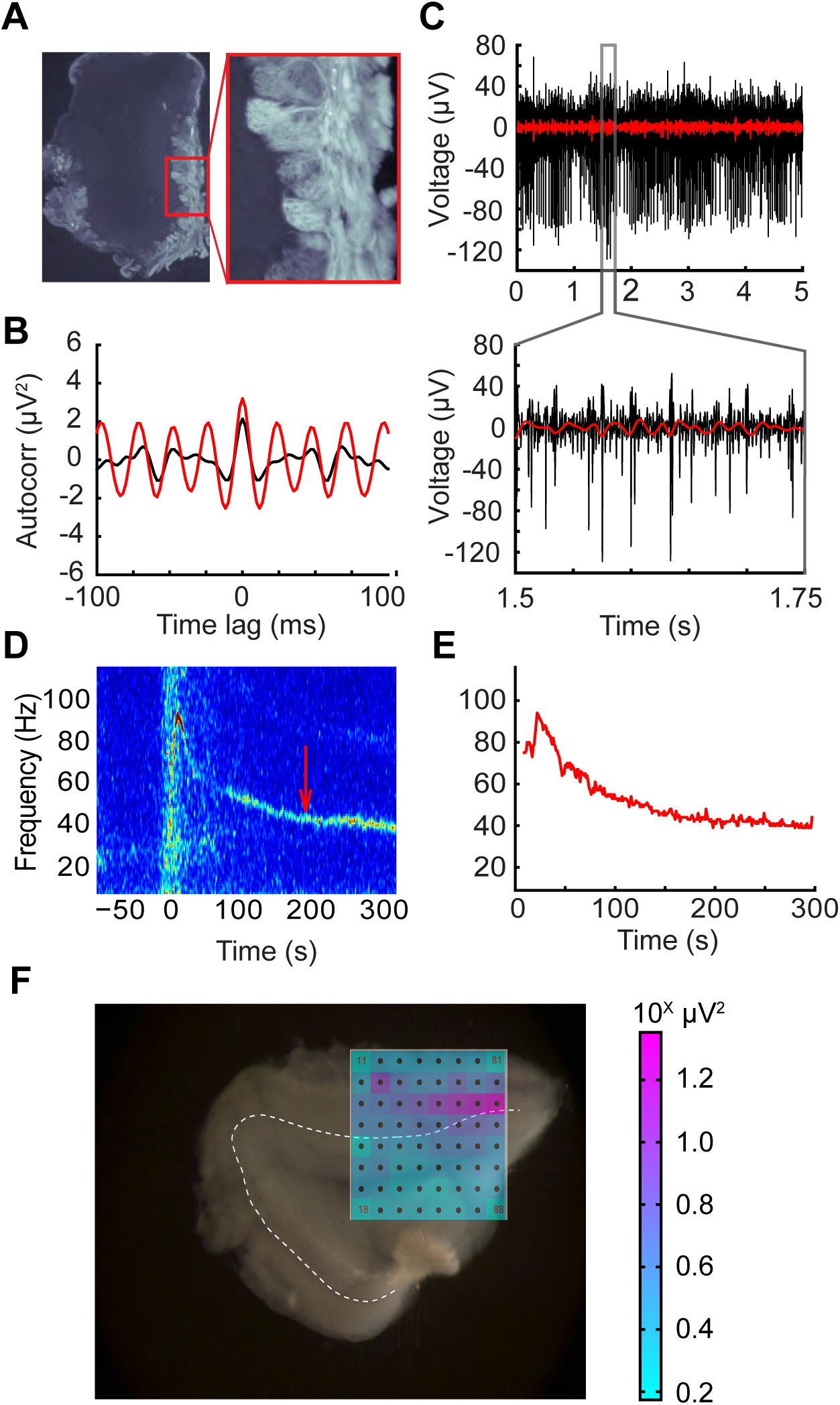
MEA recordings from olfactory bulb slices. ***A*:** 25 μm fixed OB slice from an OMP-ChR2-EYFP plasmid transgenic mouse (ORC-M) imaged with a fluorescent microscope. ChR2-EYFP coexpression in OSN axonal arbors is limited to the glomerular layer. *Inset* (20x magnification) shows distinguishable glomeruli. ***B*:** Autocorrelation of 250 ms of bandpass-filtered data (20-70 Hz; *red*) over three minutes following stimulation with 100 μM of the group I/II mGluR agonist ACPD, compared to baseline activity 50 sec prior to agonist application (*black*). Transient ACPD presentation generated a strong ∼40 Hz gamma oscillation in the slice. ***C*:** Single electrode recording from spontaneously firing presumptive MC used to align slice to array. *Lower panel* depicts a 250 ms window revealing the presence of multiple spike waveforms. Red overlay shows bandpass filtered data in the OB “slice gamma” range (20-70 Hz). ***D*:** Single-electrode spectrogram illustrating an ACPD-induced persistent gamma oscillation. The ACPD agonist was delivered at time = 0. *Arrow* indicates the starting time of the data window analyzed in panel C. ***E*:** Automated gamma oscillation detection (see *Methods*) extracts high-power regions of oscillations for array-wide visualization and further analysis. Peak extractions are from spectrogram data depicted in panel D. ***F*:** MEA schematic overlaid across an OB slice. Interelectrode distance is 200 µm. Schematic depicts gamma-band (20-70 Hz) LFP power in each electrode integrated across 5 seconds of recording following stimulation with ACPD (pink saturation indicates higher gamma-band power in log scale; i.e., units are in powers of 10). The majority of gamma band activity occurs in the external plexiform layer (EPL) immediately superficial to the mitral cell layer (*dotted line*).

### OB slice preparation for array recordings

Horizontal slices (300 μm) were prepared from the olfactory bulbs of 28-42 day old mice. Mice were anesthetized with 2% isoflurane and ketamine (150 mg/kg ip), then decapitated, after which the olfactory bulbs were quickly removed. Slices were cut on a vibrating microtome (Integraslice 7550 PSDS or Leica VT1000S) in an ice cold, oxygenated (with carbogen: 95% O2, 5% CO2), low-calcium/high-magnesium artificial cerebrospinal fluid (aCSF) dissection solution containing (in mM): NaCl 124, KCl 2.54, NaHCO3 26, NaH2PO4 1.23, CaCl2 1, MgSO4 3, glucose 10 (Balu & Strowbridge 2007). Slices then were incubated in oxygenated dissection solution at 37°C for twenty minutes, and then removed from incubation and maintained in this solution at room temperature until transfer to the recording well.

### Electrophysiological array recordings

During recordings, slices were continuously superfused at 1 ml/minute with heated (34°C), oxygenated aCSF (Gire & Schoppa 2008) (in mM: NaCl 125, KCl 3, NaHCO3 25, NaH2PO4 1.25, CaCl2 2, MgCl2 1, glucose 25) from a gravity-fed perfusion system. Slices were held in position by nylon webbing glued to a C-shaped chrome wire weight. Extracellular signals were recorded from OB slices using a 60-electrode planar microelectrode array (MEA; Multichannel Systems, Reutlingen, Germany). The array’s 60 titanium nitride electrodes (30 µm width, 30-50 kΩ impedance) were arranged in an 8x8 grid (less the four corners) at a 200 µm pitch. Electrode 15 was used as a reference. Each electrode on the MEA chip was individually connected to an amplifier in the MEA1060 baseplate. Signals were bandpass filtered (1-3000 Hz), amplified (1200x gain), sampled (5-20 kHz at 16-bit resolution), and acquired using the vendor’s MC_RACK software.

### Pharmacology

Slices were chemically stimulated by pipetting 100 ul of agonist directly onto the slice in the recording well. Both the cholinergic agonist carbachol (CCh; 186 μM) and the group I/II metabotropic glutamate receptor agonist 1-amino-1,3-dicarboxycyclopentane (ACPD; 100 μM) were able to evoke gamma oscillations (Whittington et al 1995). The group I mGluR agonist 3,5-dihydroxyphenylglycine (DHPG; 125 μM) also was tested, and produced responses similar to ACPD (not shown). Notably, these agonists diluted rapidly (assessed by dye-pipetting studies; not shown), indicating that the oscillogenic effects of these agents long outlasted their presence in the bath. Delivery of plain aCSF by pipet evoked no physiological responses, indicating that temperature or mechanical perturbations were not responsible for the observed effects. The GABA(A) receptor antagonists bicuculline methiodide (BMI) and gabazine, and the GABA(A) receptor agonist muscimol, were bath-applied. We used BMI at various concentrations in our initial efforts to ensure that GABA(A) receptors were fully blocked; all concentrations used yielded similar results and were pooled for analysis. Most GABA(A) receptor blocking experiments used 50 μM BMI. All pharmacological agents were purchased from Sigma-Aldrich (St. Louis, MO, USA).

### Optical stimulation for MEA recordings

A light engine (Lumencor, Beaverton, OR, USA) coupled to a light guide was used to optically stimulate OB slices taken from OMP-ChR2-EYFP transgenic mice. Slices were stimulated with full-field 4 Hz tetanic bursts of blue light (50% duty cycle; 15 mW/mm^2^ at 475 nm) for 5 seconds.

### OB slice preparation for intracellular recording

Horizontal slices (300 μm) were prepared from the olfactory bulbs of 28-63 day old mice. Mice were anesthetized with isoflurane and decapitated, after which the calotte was lifted and the brain was gently removed. Slices were cut on a Microm HM650V slicer (Sigmann Elektronik, Hüffenhardt, Germany) in an ice-cold, oxygenated (95% O2, 5% CO2), low-calcium/high-magnesium aCSF solution containing (in mM) NaCl 125, KCl 2.5, NaHCO3 25, NaH2PO4 1.25, CaCl2 1, MgCl2 2, glucose 25, adjusted to 310 - 320 mOsm and pH 7.2 - 7.3. Slices then were incubated in standard (low-magnesium) aCSF (in mM): NaCl 125, KCl 2.5, NaHCO3 25, NaH2PO4 1.25, MgCl2 1, CaCl2 2, glucose 25, adjusted to 310 - 320 mOsm, and bubbled with carbogen (95% O2, 5% CO2) to stabilize the pH at 7.2 - 7.3. Prior to electrophysiological recordings, slices were first incubated at 37°C for 30 min and then at room temperature for another 30 min. With the exception of glucose (Sigma-Aldrich, St Louis, USA), all chemicals listed were purchased from Merck (Darmstadt, Germany).

### Intracellular recordings

Slices were transferred to a recording chamber, and continuously superfused with heated (33-35°C), oxygenated, low-magnesium aCSF maintained by a vacuum pump (WISA, ASF THOMAS Industries GmbH, Puchheim, Germany). Whole cell current clamp recordings were performed using borosilicate glass tubing pulled to 5 - 10 MΩ resistance. Patch pipettes were filled with solution containing (in mM): KMeSO4 130, HEPES 10, KCl 7, ATP-Na 2, ATP-Mg 2, GTP 0.5, EGTA 0.05, biocytin 10, with osmolarity adjusted to 290 - 295 mOsm and pH to 7.4 (with KOH). Alexa Fluor 594 hydrazide (20 mM, Invitrogen, Carlsbad, USA) also was added to enable fluorescence imaging of the patched cell. Recordings were amplified with an Axon MultiClamp 700B microelectrode amplifier (Molecular Devices, Sunnyvale, USA) and digitized at 3 kHz. Measured potentials were not corrected for junction potentials. All chemicals were purchased from Sigma-Aldrich (St Louis, MO, USA) except for HEPES (GERBU Biotechnik GmbH, Wieblingen, Germany), EGTA (Carl Roth GmbH, Karlsruhe, Germany) and KCl (Merck, Darmstadt, Germany).

Slices were visualized using a custom-built upright two-photon microscope (Denk et al., 1990) based on a diode-pumped solid-state Verdi Laser System (10 Watt, ∼900 nm; Coherent, Santa Clara, USA). A water immersion Olympus 20x / 0.95 NA XLUMPlanFl objective (Olympus, Melville, NY, USA) was connected to the microscope and optically coupled to the tissue. Laser scanning was controlled by custom scanning software (CFNT; Ray Stepnoski, Bell Labs and Michael Mueller, Max Planck Institute for Medical Research and Lucent Technologies). Two photosensor modules (Hamamatsu Photonics K. K., Shizuoka-ken, Japan) collected the emitted light selected by two different band-pass filters (EYFP: HQ 535/30; Alexa 594: HQ 645/75; both oriented at 18°) to monitor the expressed EYFP in OSNs and the intracellular Alexa Fluor 594 hydrazide signals. One filter was mounted to one photosensor. The scanned frame size ranged from 128 x 128 pixels to 512 pixels x 512 pixels (1 - 2 ms per line).

### Optical stimulation in vitro

For spatiotemporally controlled patterns of light delivery, we coupled a DLP projector (EX330e, Optoma, Hertfordshire, UK) into the light path. To minimize vibrations, we replaced the internal cooling fan with a custom-built ventilation shaft. The projector light was directed to a movable mirror between the tube lens and the scan lens, enabling us to switch between imaging and illumination. In front of the projector we placed two lenses. One objective lens (1.6x, WD 55 mm, Leica Microsystems, Wetzlar, Germany) was optically coupled to an achromatic doublet lens (200 mm focal length; Spindler & Hoyer, Crownhill, UK). An air immersion Olympus XL Fluor 4x / 340 (0.28 NA) objective (Olympus, Melville, NY, USA) was appended to the end of the light path, above the tissue plane.

To control the timing of the light presentation, a PRONTOR magnetic shutter (PRONTOR GmbH, Bad Wildbad, Germany) was positioned between the doublet lens and the movable mirror. The shutter opening and closing was triggered by a TTL signal controlled with IGOR PRO software (WaveMetrics, Lake Oswego, OR, USA) using Neuromatic (www.thinkrandom.com). The rectangular shapes and intensity of the light stimuli were adjusted using Microsoft Powerpoint or custom-written algorithms in MATLAB (Mathworks, Natick, MA, USA). The duration of light stimuli was 100 ms. To calibrate the timing of light stimulus delivery, a light sensitive diode was connected to an LT1806 operational amplifier and positioned at the focal plane. The light onset delay following the TTL signal was measured using a custom-written algorithm in MATLAB.

### Surgery and in vivo intracellular recordings

In vivo recordings were performed as previously described (Fukunaga et al., 2014). In brief, animals were anesthetized intraperitoneally with ketamine and xylazine (100 mg/kg and 20 mg/kg, respectively for induction; xylazine concentration was reduced to 10 mg/kg for maintenance) and kept warm (37°C; DC temperature controller, FHC, Bowdoin, ME, USA) for the duration of the experiments. A craniotomy and durectomy of approximately 1000 μm in diameter was made over the dorsal part of the left olfactory bulb, which was submerged in Ringer’s solution containing (in mM): NaCl (135), KCl (5.4), HEPES (5), MgCl2 (1), CaCl2 (1.8), pH adjusted to 7.2, and 280 mOsm/kg. Whole-cell recordings were made with borosilicate glass pipette filled with (in mM): KMeSO4 (130), HEPES (10), KCl (7), ATP2-Na (2), ATP-Mg (2), GTP (0.5), EGTA (0.05), biocytin (10), with pH and osmolarity adjusted to 7.3 and 275-80 mOsm/kg, respectively. Signals were amplified and filtered at 30 kHz by an Axoclamp 2B (Molecular Devices, Sunnyvale, CA, USA) and digitized at 20 kHz with a micro1401 (Cambridge Electronic Design, Cambridge, UK). Projection neurons (mitral / tufted cells; *n* = 3) and interneurons (granule cells; *n* = 6) were identified based on depth, passive membrane properties and AHP (Kollo et al., 2014) No distinction was made between mitral and tufted cells, collectively referred to as MCs.

### Optical stimulation in vivo

For spatiotemporally controlled patterns of light delivery, the same DLP projector coupled into the light-path of a custom-built two-photon microscope was used as above and controlled with custom-written Matlab scripts. Light spots of 267 µm diameter were delivered in a grid around the recording electrode. The position of maximal depolarization was determined and further refined by applying smaller (133 µm) spots. For assessing the distance-dependence of light-evoked excitation and inhibition, concentric circles of light (100 µm thick, diameters varied) were presented, each centered around the spot of maximal activation. Whereas the distance, and not the total integrated intensity of illumination, was the independent variable of interest, stimuli with larger diameters did deliver more total light to the brain, which we examined separately (Supplementary Fig. S1).

### Spectral processing

MEA recordings were resampled at 512 Hz and analyzed in MATLAB. Principal component analysis (PCA) was used to denoise array-wide resampled datasets to remove 60 Hz interference and other highly correlated noise sources (Fig. 2b,d). PCA was computed via the covariance method. The covariance matrix **C** is the normalized covariance of the data matrix **X** comprised of all array channels. The matrix of eigenvectors **V** is computed such that **V^-1^CV** = **D**, where **D** is the diagonal matrix of eigenvalues of **C**. The principal component is removed to create a denoised data matrix **X_d_** via **X_d_** = **X** - **Xvv^T^**, where vector **v** is a feature vector comprised of the eigenvector with the highest eigenvalue.

**Figure 2.**
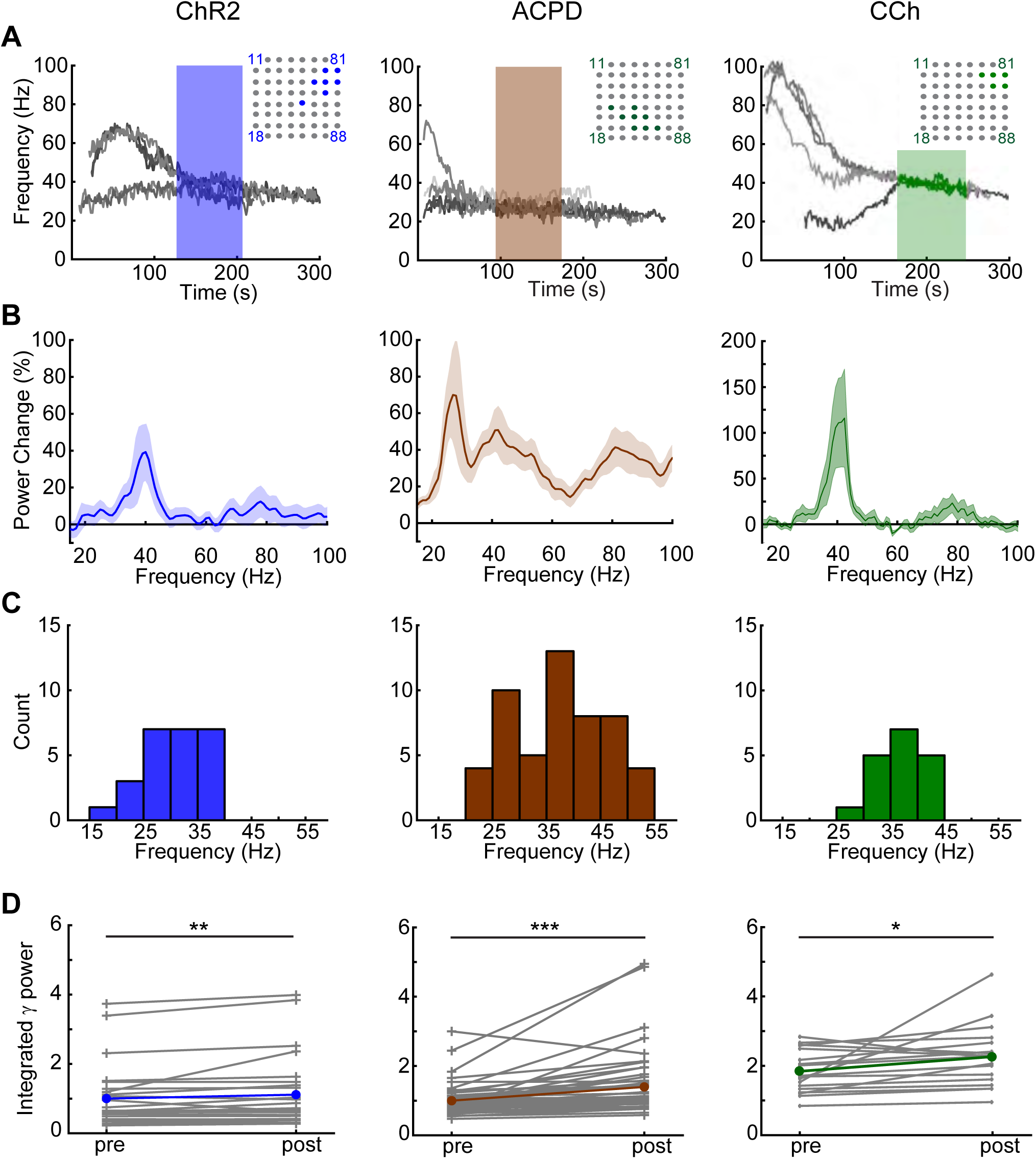
Induced gamma oscillations recorded across multiple electrodes. ***A*:** *Left panel*. Overlaid recordings from multiple MEA electrodes on a single slice following ChR2 stimulation with blue light. Seven electrodes exhibited robust gamma oscillations and were included in analyses; their locations are highlighted within the MEA schematic (*upper right inset*). The highlighted segment of the gamma traces denotes the 80-second quasi-steady-state oscillation data used for all data analyses. Note that the seven oscillations in this particular slice converge onto two stable gamma frequencies rather than one (see text and Fig. 4A*-C*). *Middle panel*. Overlaid recordings from eight MEA electrodes on a single slice simultaneously exhibiting gamma oscillations following application of 100 ul of the mGluR I/II agonist ACPD (100 μM). *Right panel*. Overlaid recordings from five MEA electrodes on a single slice simultaneously exhibiting gamma oscillations following application of 100 ul of the cholinergic agonist carbachol (186 μM). Note that the induced oscillations persist long after the termination of optical stimulation or the estimated washout times. ***B*:** Overlaid average power spectra following ChR2 activation (*n* = 11 slices/25 electrodes), ACPD application (n = 11 slices/53 electrodes), or CCh application (*n* = 5 slices/18 electrodes), with emergent power normalized to baseline spectral power at each frequency (see *Methods*) such that the plots illustrate the changes in neural activity spectra arising from the effects of each stimulation. Shading depicts the SEM. ***C*:** Histogram showing the average peak frequencies from each 80 s traced oscillation induced by ChR2 stimulation (25 electrodes), ACPD (53 electrodes), or CCh (18 electrodes). The multiple peaks shown in the ACPD power spectrum reflect variance among slices more than genuine multimodality; among ACPD-activated slices, 36 of 53 electrodes exhibited a single spectral peak, whereas the remainder exhibited two peaks or broadband activation. ***D*:** Integrated gamma-band power in each individual electrode (*grey lines*) and the means across electrodes (*colored lines*), both before (*pre*) and after (*post*) the stimulation of ChR2 or the application of ACPD or CCh. *Asterisks* reflect statistical significance; all three methods of stimulation induced significant increases in integrated gamma band power compared to baseline activity (Wilcoxon signed rank tests; *ChR2*, T(*n*=25) = 60, *p* < 0.01; *ACPD*, T(*n*=53) = 48, *p* < 0.001; *CCh*, T(*n*=18) = 31, *p* < 0.02).

Spectrograms were computed with fast Fourier transforms (FFT) using 1 second time intervals convolved with a triangular window. Intervals overlapped by 50% (i.e., 500 ms). The spectrograms were smoothed using a two-dimensional Hamming window (16 points, 4 Hz by 4 sec). To extract LFP time series for further analysis, recordings were bandpass filtered with a 2^nd^ order Butterworth filter (20Hz ≤ *f_BP_* ≤ 70 Hz). Biased autocorrelations were performed on 250 ms windows of bandpass-filtered data (Fig. 1*B*).

### Detection and extraction of gamma oscillations

The “slice gamma” band (20–70 Hz) extends to a lower frequency range than is customary for the gamma band *in vivo* (Brea et al 2009, Friedman & Strowbridge 2003, Gire & Schoppa 2008). Oscillations across this full range are considered to reflect gamma within OB slices for functional reasons; *in vivo*, gamma oscillations are endogenous to the OB whereas functionally distinct beta-band oscillations in the OB require interactions with piriform cortex and would therefore not be observed in the isolated OB slice. To isolate persistent gamma oscillations induced in OB slices, we first thresholded raw spectrograms (i.e., prior to smoothing via the Hamming window) at two standard deviations above the mean gamma band power, where each pixel represented 1 second time and 1 Hz bandwidth. This stringent criterion typically left small gaps in the gamma trace between clusters of putative gamma activity. These clusters were bridged via 1 standard deviation pixels, after which sufficiently large clusters (>5 sec) were identified as possible gamma oscillations. These clusters then were used as a mask on raw spectrograms for pathfinding. Beginning and end points in time were manually selected on the spectrogram, and the path between these points then was automatically plotted along the highest power pixels within the mask. Identified oscillations that exhibited a poststimulation shift in the peak power of the FFT and a stable peak frequency (<10 Hz variation for 80 sec; analysis window) were selected for analysis. When oscillatory frequencies are reported as X ± Y Hz, the uncertainty Y is the standard error of the mean.

### Analyses of power spectra

The frequency of LFP gamma oscillations within a given recording was determined by the mean peak frequency across an 80 sec analysis window (Fig. 2*C*). To assess the effects of optical stimulation or pharmacological manipulations on the integrated gamma band power, we compared 80 sec analysis windows from before and after the manipulation. The gamma power (*P_γ_*) was calculated between 20 and 55 Hz (Fig. 2*C*) by

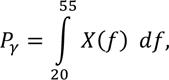

where *X(f)* is the FFT of the 80 sec analysis window. Statistical significance was assessed using Student’s t-test on the pre- and post-stimulus integrated gamma powers. Overlaid power spectra were computed with the same analysis windows using Welch’s power spectral density estimate, after which the post-stimulus density (*P_post_*) at frequency *f* was normalized to the pre-stimulus density (*P_pre_*):

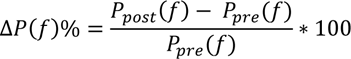

yielding *ΔP(f)%,* which is the percentage change in power density generated by the experimental manipulation (Fig. 2*B*). The shaded areas of Fig. 2*B* show the standard error of the mean of *ΔP(f)%*.

*Analyses of coherence.* Coherence (*C_xy_*) between oscillations on two different electrodes was computed in the frequency domain via

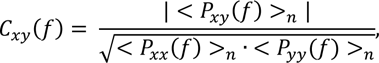

where *P_xx_* and *P_yy_* are autocorrelations and *P_xy_* is the cross-correlation between the two electrodes, each averaged over *n* 1-sec epochs (here, *n* = 80). Note that C_xy_ is a complex number from which a magnitude and phase can be derived. Coherence analyses were performed in slices that had at least four adjacent electrodes with detectable, stable gamma band oscillations persisting for at least 80 sec and a coherence magnitude (|C_xy_|) of at least 0.15 between any one such electrode and at least three neighboring electrodes. The frequency with the highest coherence magnitude (Fig. 4*A-B*) was selected as the peak frequency for that coherence group (of electrodes). For a given peak frequency, the electrode that had the greatest coherence magnitude with the largest number of neighboring electrodes was designated as the *reference electrode* for that coherence group (not to be confused with the electrophysiological reference electrode, which was always MEA channel 15). Pairwise coherence spectra then were calculated between each reference electrode and all other active electrodes from a given slice, and used to produce quiver plots (Fig. 4*C*) and to calculate the average coherence across the spatial extent of the OB circuit (Fig. 4*D-F*).

### Statistical analyses of regional coherence

To assess the functional significance of coherence reductions observed under GABA(A) receptor antagonists, we first sought to distinguish ‘true’ coherence arising from the synaptic coupling of neighboring OB columns from ‘spurious’ coherence arising simply from increased levels of nonspecific neuronal activity (Buzsaki & Schomburg 2015). To do this, we estimated the baseline level of spurious coherence under each treatment condition by scrambling the pairings of reference electrodes with their corresponding peak frequencies. Pairwise coherence measurements then were made between each reference electrode and all other electrodes in the slice, producing a set of coherence magnitudes identical to those of the nonscrambled condition but no longer constrained with respect to distance. This scrambling process therefore disrupts coherence arising from correlated activity among neighbors while preserving spurious coherence arising from nonspecific activity. Subtracting this spurious coherence estimate from the total measured coherence provided a measure of the ‘true’ coherence arising from correlated activity between neighboring electrode recordings. To compare the (true) coherence evoked under aCSF to that under BMI, we estimated the true coherence between all pairs of electrodes under each condition by subtracting scrambled from unscrambled raw coherence.

### Computational modeling

A biophysically-based network model of the olfactory bulb was constructed from 64 mitral cells (MCs), 64 periglomerular cells (PGs) and 256 granule cells (GCs). All cellular and synaptic models were identical to those described previously (Li & Cleland 2017) except as specifically described below. Briefly, all neurons were oligocompartmental with 4-7 distinct membrane conductances in each compartment; individual MCs exhibited intrinsic subthreshold membrane potential oscillations as has been experimentally reported (Balu et al 2004, Chen & Shepherd 1997, Desmaisons et al 1999). Model neurons were deployed within a two-dimensional 1000 μm х 1000 μm grid; neurons of each individual type (MC, PG, GC) were arranged in square arrays with equal separation in the horizontal and vertical directions (Fig. 7*A*). In contrast to previous versions of this model (Li & Cleland 2013), the connection probability between MCs and GCs was distance-dependent according to

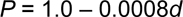

where *d* is the distance in μm between their somata on the surface; i.e., connection probabilities declined linearly from 1.0 when adjacent to 0.2 at 1000 μm distance. To model the slice preparation, in which longer lateral projections are increasingly likely to be disrupted, connectivity with a given MC followed the same distance function but was restricted to GCs within a 250 μm radius of its soma (Fig. 7*A*, *circle*). Steady-state odor inputs were drawn randomly from a uniform distribution across [0.2, 1.0]. LFP responses to simulated odorant presentations then were measured by averaging and filtering MC membrane potential fluctuations across the layer, simulating recordings obtained by a single electrode sampling a large field (see *Discussion*; also see Li and Cleland (2013) for details). The GABA(A) decay time constant for GC-to-MC synapses was 3 ms (the effects of this time constant are explored in Litaudon et al. (2008)).

## Results

### Persistent gamma oscillations are evoked by optical stimulation of sensory neuron arbors or application of chemical agonists

Horizontal slices of mouse olfactory bulb were laid onto a 60-channel planar microelectrode array (MEA; Multichannel Systems, Reutlingen, Germany) and aligned with the aid of endogenous spiking activity measured from the large projection neurons of the deep EPL and mitral cell layers. No spiking activity was observed under these conditions from the glomerular layer or from layers deep to the mitral cell layer. Local field potentials recorded from the external plexiform layer (EPL) under control conditions in aCSF occasionally exhibited spectral peaks in the 20-70 Hz “slice gamma” band (henceforth *gamma band*; see *Methods*), but usually exhibited no stable peak frequencies prior to stimulation. Either optical stimulation of channelrhodopsin-2 (ChR2)-expressing OSN axonal arbors within the glomerular layer (4 Hz for 5 seconds, 50% duty cycle; Fig. 1*A*) or transient application of either the Group I/II metabotropic glutamate receptor (mGluR) agonist 1-amino-1,3-dicarboxycyclopentane (ACPD) or the cholinergic agonist carbachol (CCh) induced spiking activity and persistent gamma band LFP oscillations across the EPL (Fig. 1*B-F*).

Oscillations recorded from neighboring MEA electrodes (200 μm pitch) were independent, but often converged onto common stable gamma frequencies. All spectral analyses were based on 80-second analysis windows that began once all of the spectral peaks from electrodes recording from a given slice had stabilized to consistent frequencies within a 10 Hz band (Fig. 2*A*; *shaded regions of spectrograms*). Both ChR2 optical stimulation and activation with ACPD or carbachol evoked strong spectral peaks in the gamma band compared to baseline activity (Fig. 2*B*). Histograms of the mean peak frequencies from all recordings confirmed a relatively narrow peak among ChR2-activated slices (*n* = 11 slices, 25 electrodes) and CCh-activated slices (*n* = 5 slices; 18 electrodes), and a broader set of peak frequencies in ACPD- activated slices (*n* = 11 slices, 53 electrodes; Fig. 2*C*). Among ACPD-activated slices, 36 of 53 electrodes exhibited a single peak at a frequency that ranged from 30 to 55 Hz, whereas the remainder exhibited two peaks or broadband activation. Gamma-band power was significantly increased under all of these methods of activation: optical stimulation of ChR2-expressing afferents (Wilcoxon signed-ranks test; T(*n*=25) = 60; *p* < 0.01), ACPD activation (T(*n*=53) = 48; *p* < 0.001), and CCh activation (T(*n*=18) = 31, *p* < 0.02); this increase was observable in nearly all individual recordings (Fig. 2*D*). The action potentials of a subset of active neurons – presumed to be principal neurons owing to their localization to the mitral cell layer and deep EPL – were constrained with respect to the phase of evoked gamma oscillations (Fig. 3*A*), consistent with previous findings (Bathellier et al 2006, Eeckman & Freeman 1990, Kashiwadani et al 1999, Werth et al 2022). This phase constraint occurred with all methods of activation, though it was clearest in response to ChR2 stimulation (Fig. 3*B-D*). Irrespective of the method used to generate oscillations, gamma power was restricted to the EPL (Fig. 1*F*).

**Figure 3.**
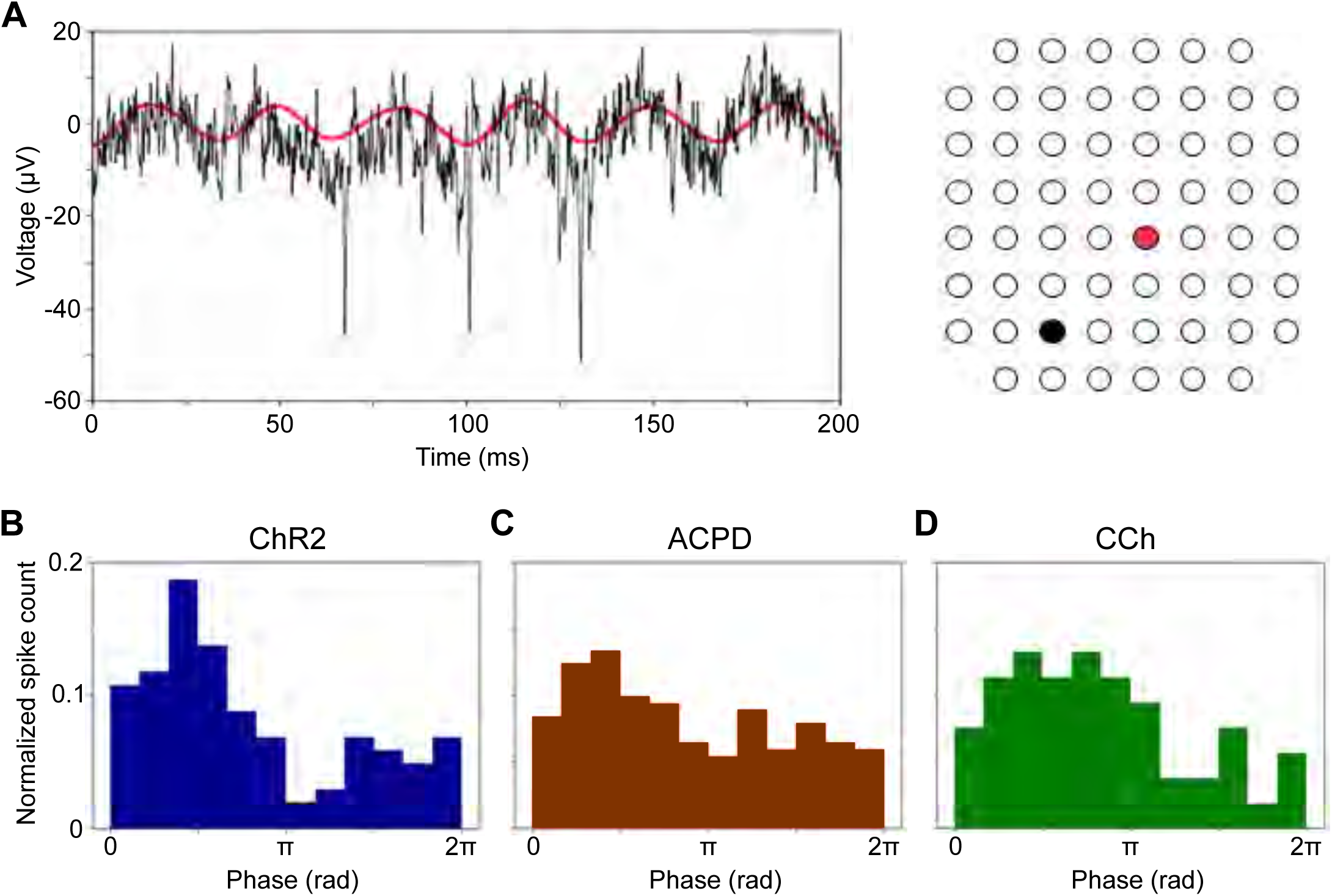
Gamma oscillations phase-constrain evoked MC action potentials. ***A*:** Unfiltered recordings from one electrode (*black trace*) show principal neuron action potentials that are phase-constrained with respect to the coherent gamma oscillations simultaneously recorded from a separate electrode (*red trace*; interelectrode distance = 565 μm). *Right panel* depicts the electrodes’ locations. ***B-D*:** Spike phase histograms for action potentials recorded from a principal neuron following (B) optical stimulation of ChR2 or chemical stimulation with (C) ACPD or (D) carbachol as in Fig. 2.

### Gamma oscillations form localized regions of coherence in slices

Evoked gamma oscillations often were coherent across multiple electrodes. To quantify the spatial patterns of coherence across the EPL, we selected each active electrode (i.e., each electrode exhibiting gamma-band activity) in turn and computed its coherence with all other active electrodes. For a given peak frequency, the electrode that had the greatest coherence magnitude with the largest number of neighboring electrodes was designated as the *reference electrode* for that coherence group. Pairwise coherence spectra then were calculated between each reference electrode and all other active electrodes from a given slice (Fig. 4*A-B*). Notably, some OB slices simultaneously exhibited multiple discrete localized patches of coherent gamma oscillations with different frequencies (Fig. 4*C*). These results suggested that groups of neighboring OB columns were synchronized through synaptic interactions, with sharp discontinuities between local regions of coherence arising from the reduced density of lateral interactions produced by the slicing process – a phenomenon theoretically predicted by two-dimensional map-based models (Bazhenov et al 2008) and illustrated below. A similar phenomenon has been observed *in vivo* under urethane anesthesia (Neville & Haberly 2003), which is thought to induce concerted, individually modest effects on multiple excitatory and inhibitory synaptic receptors (Hara & Harris 2002).

**Figure 4.**
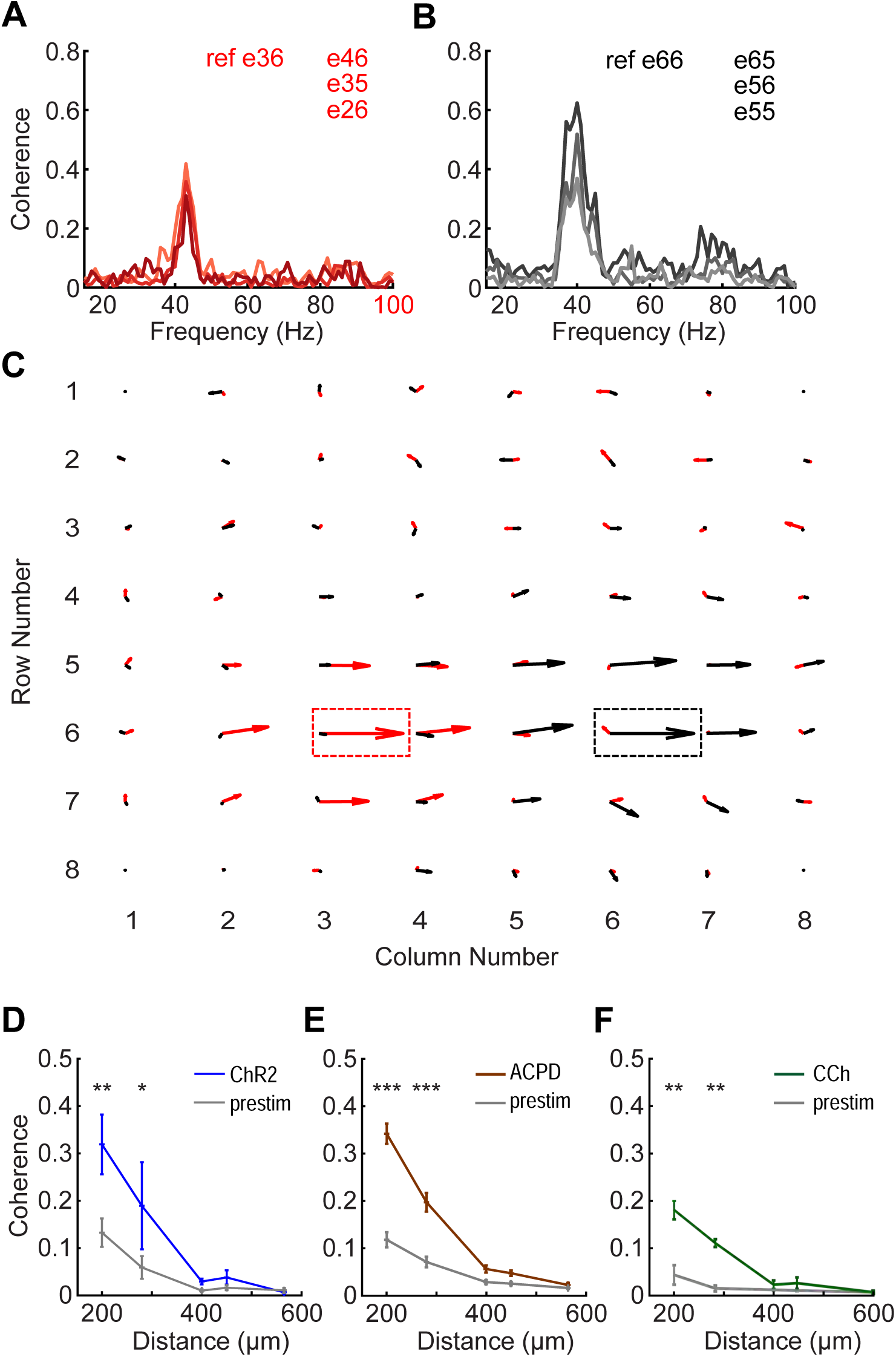
OB slices can exhibit multiple local regions of coherent gamma oscillations. ***A*:** Coherence between a selected reference electrode (e36; nomenclature based on XY coordinates as shown in panel C) and three adjacent electrodes in a selected slice. The frequency with the highest coherence magnitude in this region is 43 Hz. ***B*:** Coherence between another reference electrode (e66) and three additional adjacent electrodes, in the same slice depicted in panel A. The frequency with the highest coherence magnitude in this region is 40 Hz. ***C*:** Quiver plot illustrating these two adjacent, simultaneous regions of coherence respectively centered on reference electrodes e36 (43 Hz, *red arrows*) and e66 (40 Hz, *black arrows*). Reference electrodes are boxed. The lengths of the arrows indicate coherence magnitude and the angles indicate phase; rightward facing arrows denote zero phase lag with respect to the corresponding reference electrode. Note the nonoverlapping regions of coherence for each reference frequency. ***D*:** Plots of mean coherence magnitude versus inter-electrode distance before and after ChR2 optical stimulation. The ordinate depicts the average coherence magnitude based on 80 s of quasi-stationary data averaged across all pairs of active electrodes separated by the distance depicted on the abscissa (200-565 μm). Inter-electrode coherence was significantly increased by ChR2 stimulation at distances of 200 and 280 μm, but not at longer distances (*n* = 2 slices, 2 electrodes; 200 μm: *p* = 0.003; 280 μm: *p* = 0.026; see text for details). Error bars indicate SEM. ***E*:** The application of ACPD also significantly increased gamma-band coherence between electrodes at distances of 200 and 280 μm, but not at longer distances (*n* = 10 slices, 15 electrodes; *p* < 0.001 for both distances). ***F*:** The application of carbachol also significantly increased gamma-band coherence between electrodes at distances of 200 and 280 μm, but not at longer distances (*n* = 2 slices, 3 electrodes; *p* < 0.01 for both distances).

We measured the mean spatial extent of coherent regions in slices by averaging the peak coherence magnitudes between reference electrodes and all other electrodes (whether active or not) across all slices as a function of distance from the reference electrode. Both the optogenetic stimulation of ChR2- expressing OSN arbors (Fig. 4*D*) and the transient application of chemical agonists (Fig. 4*E-F*) generated significant increases in gamma-band coherence across moderately distant electrodes compared to prestimulation coherence. Specifically, two-factor analysis of variance (ANOVA) for ChR2 data demonstrated highly significant effects of *stimulation* (F(1,60) = 7.467, *p* < 0.01) and *distance* (F(4,60) = 11.043, *p* < 0.001); post hoc testing indicated significant effects of ChR2 stimulation on coherence between electrodes at 200 (*p* = 0.003) and 280 μm (*p* = 0.026) distances (Fig. 4*D*). A similar analysis of the ACPD data revealed highly significant effects of both *stimulation* (F(1,584) = 110.553, *p* < 0.001) and *distance* (F(4,584) = 114.463, *p* < 0.001); post hoc tests indicated significant effects of ACPD stimulation on coherence at 200 and 280 μm (*p* < 0.001 in both cases; Fig. 4*E*). Finally, analysis of the CCh data also revealed highly significant effects of both CCh *stimulation* (F(1,112) = 29.8, *p* < 0.001) and *distance* (F(4,112) = 15.77, *p* < 0.001, with post hoc testing indicating significant effects of CCh on coherence at 200 and 280 um (*p* < 0.01 in both cases; Fig. 4*F*)). Coherence differences measured at distances above 400 μm should not be interpreted as meaningful in any event, because the low absolute coherence values at these distances (roughly, below 0.1) are reasonably likely to arise for spurious or artifactual reasons (Buzsaki & Schomburg 2015).

### GABA(A) receptor blockade reduces the spatial coherence, not the power, of gamma oscillations

The observed distinct regions of coherence in OB slices provided a unique opportunity to study the construction and underlying mechanisms of spatial coherence in neuronal networks. To test whether regional coherence arose from lateral inhibitory synaptic coupling, we bath-applied GABA(A) receptor antagonists and measured the resulting changes in gamma oscillatory power and regional coherence. Surprisingly, ACPD-evoked gamma-band oscillations persisted in the presence of saturating concentrations of GABA(A) receptor antagonists (Fig. 5*A*) – a result superficially at odds with prior reports, using single LFP electrodes, that indicated a strong sensitivity of OB gamma oscillations to GABA(A) receptor blockade (Bathellier et al 2006, Lagier et al 2004) (see *Discussion*). To explore this result, we computed the average power spectra of ACPD-induced oscillations from 22 electrodes across 5 slices exposed to both control conditions and GABA(A) receptor blockade. Bath application of GABA(A) receptor antagonists did not significantly affect ACPD-induced oscillatory power, but did significantly reduce the mean peak frequency of the oscillation (from 34 ± 2.15 Hz to 29 ± 1.19 Hz; paired *t*-test; t(21) = 3.27, *p* < 0.01; Fig. 5*B*). Moreover, GABA(A) blockade also significantly reduced the true coherence among electrodes (see *Methods*). Using two-factor ANOVA, BMI *treatment*, interelectrode *distance*, and their interaction were all highly significant (*treatment*: F(1,471) = 33.616, p < 0.001; *distance*: F(4,471) = 34.934, p < 0.001; *treatment*distance*: F(4,471) = 6.699, p < 0.001); post hoc testing further indicated that the application of BMI specifically impaired true coherence at 200 and 280 µm interelectrode distances (*p* < 0.001 in both cases; Fig. 5*C*).

**Figure 5.**
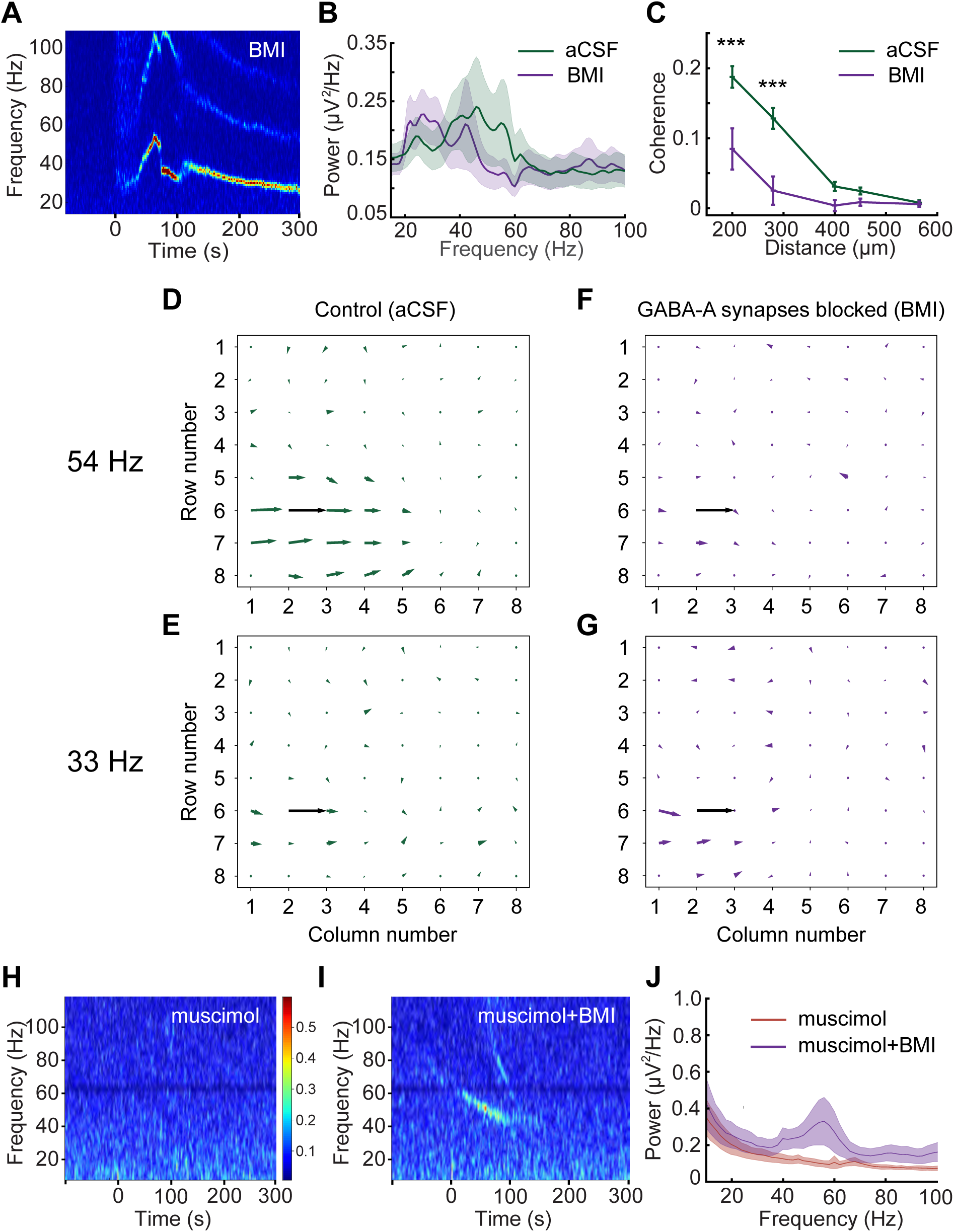
Intercolumnar synchronization is mediated by GABA(A) receptors, whereas local oscillations are GABA(A) receptor-independent. ***A*:** Spectrogram illustrating a long-lasting gamma band oscillation induced by the transient application of ACPD (100 ul bolus delivered at t=0) during bath application of 50 μM BMI. ***B*:** Average power spectra comparing oscillations induced by ACPD in an aCSF bath (*green trace*) to those induced under BMI blockade (50-500 μm BMI; *violet trace*). Only slices for which 80 s of data were recorded under both conditions were included in the analysis (*n* = 5 slices, 22 electrodes). The BMI bath induced a significant reduction in the mean oscillation frequency (34 ± 2.15 Hz to 29 ± 1.19 Hz; paired t-test; t(21) = 3.27, *p* < 0.01), but no significant change in power. Shading depicts the SEM. ***C*:** Mean true coherence with respect to distance arising from ACPD-induced oscillations in plain aCSF (vehicle control; *green*) and after the addition of the GABA(A) receptor antagonist BMI (n = 3 slices, 10 electrodes; 50-500 μM; *violet*). *Asterisks* indicate significant differences between the aCSF and BMI conditions (*p* < 0.001). ***D*:** Quiver plot illustrating the spatial extent of gamma coherence with reference electrode e26 at 54 Hz under control conditions. ***E*:** No interelectrode coherence is evident at 33 Hz under these control conditions. ***F*:** After BMI is added, interelectrode coherence at 54 Hz disappears, in part because BMI treatment reduced the peak frequency of the local coherence group to 33 Hz. ***G*:** In the presence of BMI, gamma power across active electrodes remained strong, with a mean peak frequency of 33 Hz. However, interelectrode coherence was weak to negligible, indicating that BMI treatment had decoupled these local oscillators. ***H*:** Spectrogram of OB slice activity in a 50 μM bath of the GABA(A) agonist muscimol. Muscimol reduced spontaneous activity and entirely prevented the evocation of gamma band activity by ChR2 optical stimulation (5 sec, delivered from time = 0). ***I*:** The addition of 100 μM BMI to the bath (in addition to the muscimol) rescued the ability to evoke gamma oscillations by stimulation with 475 nm blue light (5 sec, from time = 0). ***J*:** Averaged power spectra of light-induced oscillations (*n* = 4 slices, 52 electrodes) in muscimol bath (*red trace*) and in muscimol + BMI (*violet trace*). Shading depicts the SEM.

Because the power in individual electrodes was not reduced, this reduction in coherence could be attributed to phase drift (decoupling) among LFP recordings from neighboring electrodes. Indeed, analyses of recordings from one of these slices (Fig. 5*D-G*) illustrate that the broad region of coherence exhibited at the peak oscillation frequency under control conditions (Fig. 5*D*) was largely eliminated in the presence of BMI (measured at the new, BMI-induced, lower peak oscillation frequency exhibited by that slice; Fig. 5*G*). No regions of coherence were observed in control analyses of the corresponding off-peak oscillation frequencies (Fig. 5*E-F*). These results indicate that GABA(A) receptor blockade decouples OB columns from one another, eliminating cross-columnar coherence and unmasking a slower, spatially localized, GABA(A) receptor-independent oscillatory mechanism that is likely to arise from MC subthreshold oscillations ((2017); see *Discussion*). Consistent with these results, the cell bodies of MCs that innervate a single glomerulus (likely corresponding to a column) are physically distributed within regions of 100-200 um radius (Ke et al 2013).

In these experiments, to ensure that GABA(A) receptors were fully blocked, we applied the GABA(A) receptor antagonists BMI (50-500 μM) or gabazine (500 μM; data not shown) at high concentrations (compare to blockade of gamma oscillations in hippocampal slices using 1-4 μM BMI (Pais et al 2003)), and sometimes preincubated slices in the antagonist to ensure that the antagonists fully permeated the slice. However, it remained a possibility that the persistent oscillations we observed were a result of incomplete GABA(A) receptor blockade. To rule this out, we bath-applied the GABA(A) receptor agonist muscimol (50 μM), which broadly activated GABA(A) receptors, silencing all activity in the OB slice and preventing activation by ChR2 stimulation (Fig. 5*H*). Adding 100 μM BMI to the bath (in the continued presence of muscimol) rescued the excitability of the slice and restored its capacity to generate persistent gamma oscillations following ChR2 activation (*n* = 4 slices, 52 electrodes; Fig. 5*I-J*). These results indicate that localized (within-column) LFP oscillations are genuinely GABA(A) receptor-independent and inducible by afferent activation, and that these local oscillators can be coupled across multiple neighboring OB columns via GABA(A)-ergic synapses to generate broadened, inter-columnar regions of coherent activity in the gamma band.

### EPL connectivity in OB slices and in vivo

How broadly can GABA(A)-ergic synaptic connections maintain gamma-band coherence across the OB? Early studies of OB dynamics *in vivo* suggested that gamma oscillations were coherent across the entire OB (Freeman 1978); subsequent studies have largely corroborated this principle within individual respiration cycles (Kay & Lazzara 2010), but the underlying mechanisms are unclear. Maintaining global coherence across a spatially distributed network is nontrivial, and depends strongly on the density of synaptic interconnections (Rulkov & Bazhenov 2008), which in slices are substantially reduced in a distance-dependent manner. To assess whether the coherence properties and mechanisms elucidated in the slice preparation predict global coherence in an intact OB, we used intracellular recordings from MCs and GCs along with localized optogenetic stimulation to measure distance-dependent lateral synaptic connectivity both in the slice and *in vivo*. We then assessed the changes in spatial coherence attributable to these connectivity differences using a biophysically realistic computational model of OB circuitry.

To measure the spatial receptive fields of MCs in OB slices, spots of light (100 µm along the glomerular layer x 500 µm perpendicular to the layer) were delivered onto 9-12 different locations within the glomerular layer of OB slices from OMP-ChR2 mice while intracellular recordings were made from MCs or GCs (Fig. 6*A*). As in the MEA recordings, MCs were synaptically excited by the optogenetic activation of OSN axonal arbors. As predicted, spiking responses in MCs were observed within a limited radius (100-200 µm) of the stimulus spot, consistent with the distribution of MC cell bodies that innervate a single glomerulus (Ke et al 2013) (Fig. 6*B-C*; *red traces*). Moreover, broad-field optical stimulation of the entire GL with the exception of this narrow excitatory region failed to substantially excite these MCs (not shown), corroborating the narrow excitatory field of MCs and demonstrating the spatial precision of our optical stimulation.

**Figure 6.**
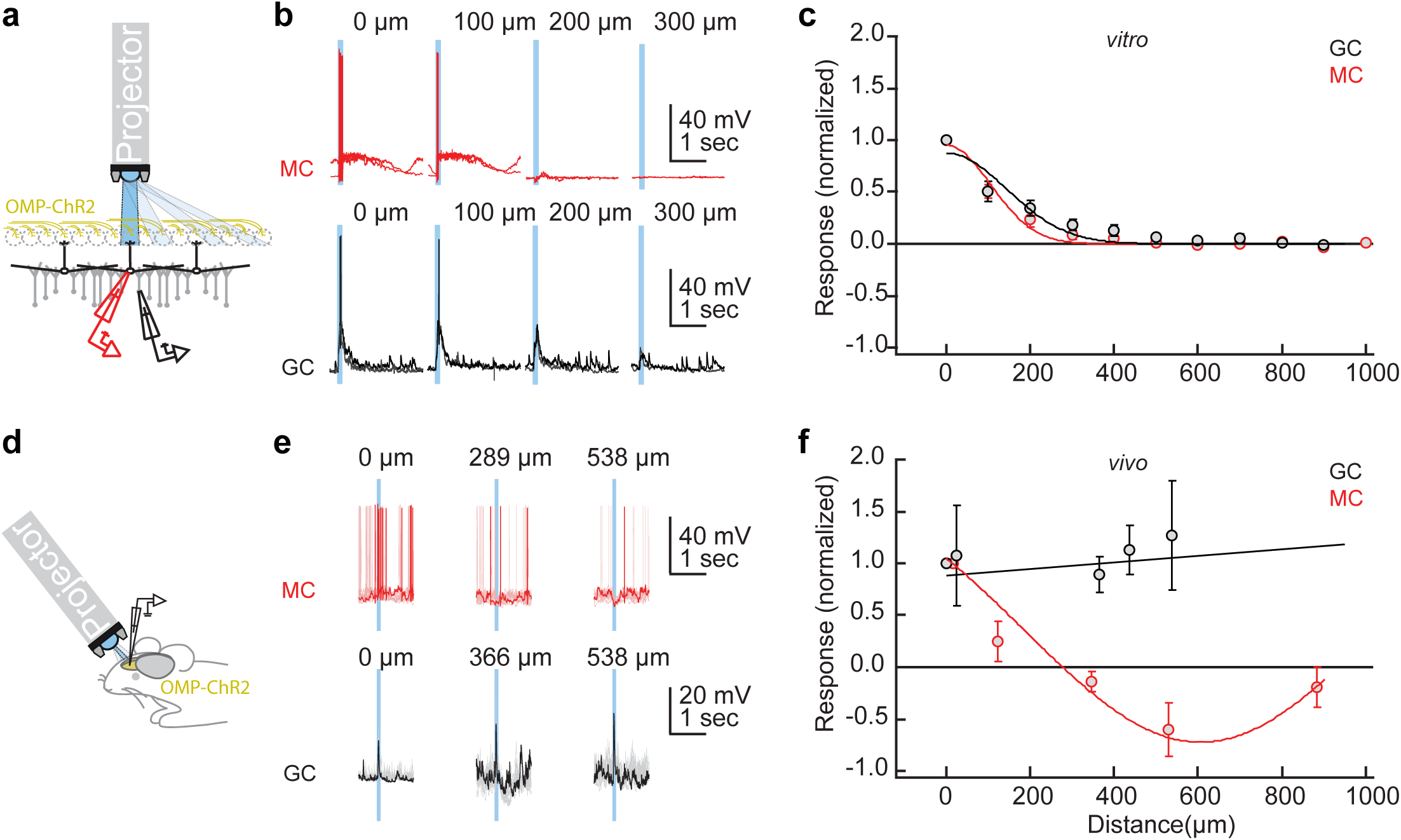
Lateral excitation and inhibition *in vitro* and *in vivo. **A-C***: *In vitro. **A***: Recording and stimulation configuration. Whole-cell patch recordings were taken from MCs (*red*) or GCs (*black*) in OB slices from OMP-ChR2 mice. A modified projector (see *Methods*) was used to illuminate 500 µm x 100 µm rectangles across the glomerular layer (GL; 500 µm along the OSN-GL-EPL axis, 100 µm along the GL). ***B*:** 100 ms blue light pulses delivered at a range of distances from the recording site evoked depolarization and action potential discharge in MCs (*red traces*) and GC (*black traces*; two repetitions each for one example cell). Distances were normalized to the illumination rectangle resulting in the strongest depolarization. ***C*:** Distance-dependence of responses (average of 0.5-1 s following the start of light stimulation) normalized to the maximum for each cell (*n* = 6-10 MCs and *n* = 6-7 GCs for distances ≤ 600 µm; *n* = 1-6 cells for larger distances). Curves are Gaussian fits. ***D-F*:** *In vivo. **D***: Recording and stimulation configuration. Whole-cell patch recordings were performed in anesthetized OMP-ChR2 mice from putative MCs and GCs from the dorsal aspect of the OB. The same projector as in (a-c) was used to deliver concentric rings of illumination (100 µm thick) centered around the region of maximal activation on the dorsal surface. ***E*:** Evoked activity in an example MC (*red trace*) and GC (*black trace*) in response to brief (100 ms) light stimulation; ten repetitions are shown with one repetition in bold. The abscissa denotes the distance from the center of the stimulus pattern to the interior diameter of the illuminated ring. Note the robust hyperpolarization of the MC at 538 µm and robust depolarization of the GC across all distances. ***F*:** Distance-dependence of responses normalized as in (c). Fitted curves are linear (for GCs; *n* = 6) and Gaussian (for MCs; *n* = 3) fits. Data in (c) and (f) are depicted as mean ± SEM.

We then similarly measured the spatial receptive fields of GCs, which receive excitation (EPSPs) from MC lateral dendrites, using the same optical stimulation protocol. Because MC lateral dendrites extend over a millimeter from the MC soma and support centrifugal action potential propagation, EPSPs in a GC could in principle arise from any MC without respect to proximity (Orona et al 1984, Shipley & Ennis 1996). However, in slices, the spatial receptive fields of GCs were relatively narrow (although somewhat broader than those of MCs), averaging 200-300 μm from the stimulus spot (Fig. 6*B-C*; *black traces*). Broad-field illumination of the entire GL except for the narrow MC excitatory region produced only very weak EPSP responses, again demonstrating the spatial precision of optical stimulation (not shown). The implication is that MCs were excited by activity within a single glomerulus, whereas GCs in the slice were excited by MC activation within a local region of a few glomerular diameters.

We then repeated these experiments in anesthetized OMP-ChR2 mice *in vivo* (Fig. 6*D*). After establishing whole-cell recordings from MCs on the dorsal aspect of the OB, light was delivered onto the dorsal surface of the OB using the same projector as in the *in vitro* experiments. To equally activate inputs from all sides (rostral, caudal, medial and lateral) of the patched MC, concentric rings (width 100 µm) were illuminated at different distances surrounding the recording site (Fig. 6*D-F*). As with the *in vitro* results, activation of MCs was reliable and efficient in close proximity but decreased rapidly with distance. Moreover, light stimulation at longer distances weakly inhibited MCs (Fig. 6*E-F*, *red traces*), consistent with the effects of long-range lateral inhibition *in vivo* (Banerjee et al 2015, Economo et al 2016, Luo & Katz 2001). Similar experiments using discrete spots of illumination (133 or 267 µm diameter) produced similar results (not shown), again demonstrating the localized excitation and broadly sourced inhibition of MCs.

As in the slice, focal illumination *in vivo* also reliably excited GCs, presumably via MC-GC synaptic activation (Fig. 6*E-F*; *black traces*). However, whereas this excitatory receptive field was confined to ≈200 µm *in vitro* (Fig. 6*C*), illumination at all distances measured from the recorded GC (as far as 500 µm) resulted in robust excitation *in vivo* (Fig. 6*F*). This indicates that lateral connections in the intact OB support the long-distance lateral excitation of GCs, thus providing a potential substrate for coordinating activity across the full extent of the OB EPL.

### Quantitative effects of lateral connectivity on coherence field

To support global coherence across the OB EPL, the extensive, spiking lateral dendrites of intact MCs should interconnect and coordinate local columnar oscillators at all distances via the mechanisms elucidated in OB slice experiments, resulting in globally coherent activity among activated neurons across the OB (Li & Cleland 2017). To demonstrate this, we modified a biophysically realistic dynamical computational model of the OB network (Li & Cleland 2013) such that cellular locations within the two-dimensional external plexiform layer were explicit, spike propagation delays were appropriate, and the density of reciprocal synaptic interactions was a declining function of physical distance within this layer (see *Methods*). To model the transected lateral dendrites of the slice preparation, connectivity with any given MC was restricted to GCs within a 250 μm radius of its soma (Fig. 7*A*, *circle*). Global LFP responses to simulated odorant presentations then were measured by averaging and filtering MC membrane potential fluctuations across the entire layer, simulating recordings obtained by a single electrode sampling a large field (Li & Cleland 2017, Viswam et al 2019).

**Figure 7.**
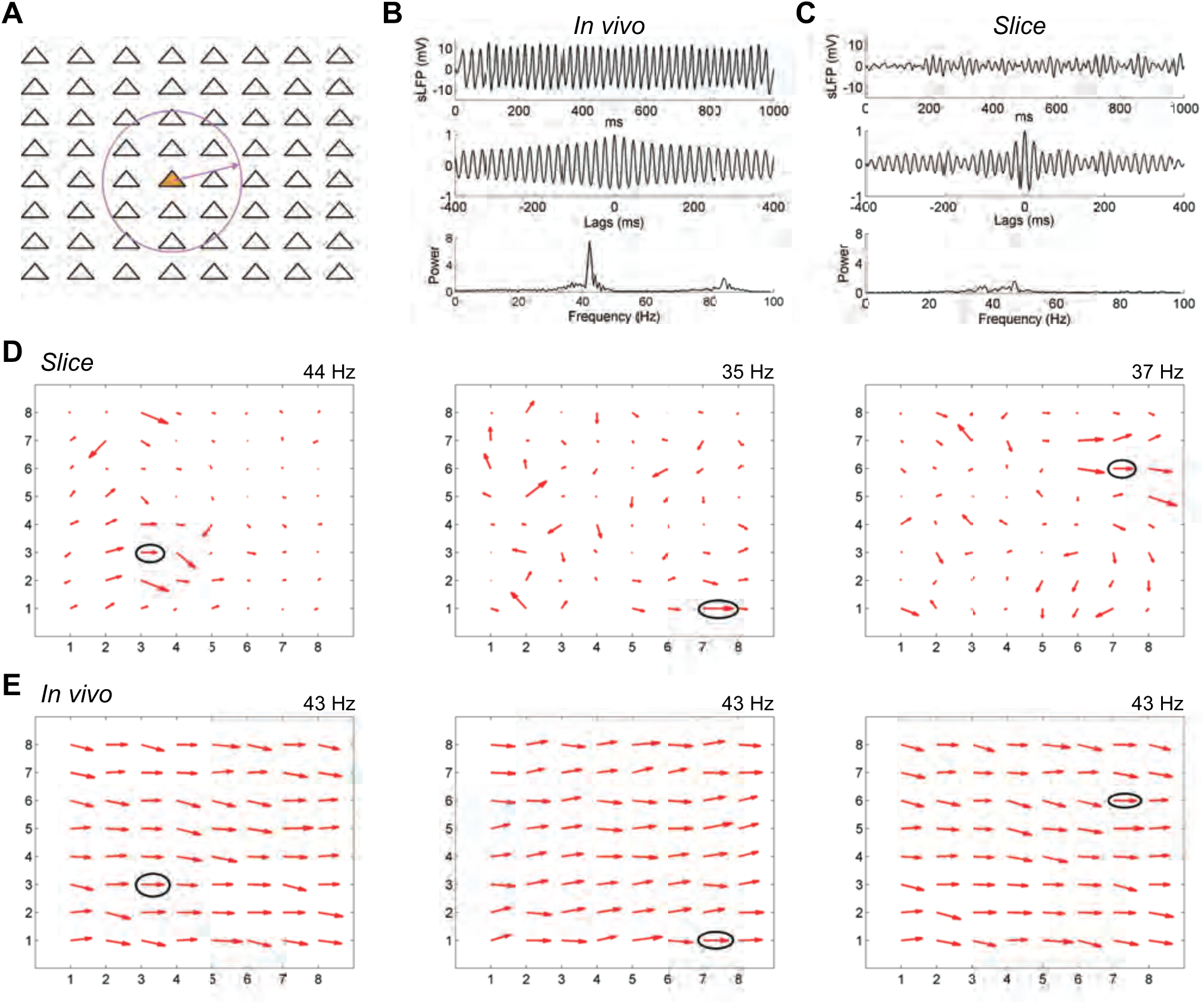
Computational modeling shows how local coherence regions in slices predict global gamma coherence in the intact OB. ***A*:** The 64 mitral cells in the model were arranged in an 8 by 8 square array with equal separation in both the horizontal and vertical directions across a two-dimensional surface (1000 μm х 1000 μm). Periglomerular cells corresponded 1:1 with MCs, whereas 256 GCs were deployed in a separate 16x16 array (not shown) across the same surface. The network connection probability between MCs and GCs declined linearly with distance in the intact (*in vivo*) OB condition. In the slice condition, the connection probability between MCs and GCs separated by greater than 250 μm (indicated by a *circle*) was reduced to zero, but was unaffected at distances of 250 μm or less. See text and Methods for details. ***B*:** Global properties of the bulbar LFP in the intact (*in vivo*) OB network model. The *top panel* depicts the simulated LFP averaged across the full network during odor presentation, the *middle panel* depicts the autocorrelation of this simulated LFP, and the *bottom panel* depicts the power spectrum. ***C*:** Global properties of the bulbar LFP in the OB network model under slice conditions. Panels are as described in B. ***D*:** Quiver plots of network coherence at three frequencies identified and measured in the slice condition. The reference electrode for each frequency is *circled*; the magnitude of coherence of each electrode with the reference electrode is denoted by the length of the corresponding arrows whereas phase with respect to the reference electrode is denoted by their angle. *Left panel* depicts the coherence profile across the electrode array at 44 Hz, *center panel* depicts the coherence profile at 35 Hz, and *right panel* depicts the coherence profile at 37 Hz. See text for details. ***E*:** Quiver plots of network coherence in the *in vivo* OB network with respect to the same three reference electrodes identified in the slice condition and depicted in (d). Note that the intact network, with its greater connectivity but otherwise identical properties, imposes a single common frequency (43 Hz) and shared phase upon the LFPs measured at all electrode sites, overpowering the preferred frequencies of local regions that were observable in the reduced coupling density of the slice condition.

In the intact (*in vivo*) model, odorant presentation generated strong global gamma oscillations (Fig. 7*B*, *top panel*) that were stable over time (*middle panel*) and exhibited a sharp spectral peak at 43 Hz (*lower panel*). In contrast, the slice model generated weak global oscillations with negligible continuity and no clear peak (Fig. 7*C*), indicating that neuronal activity was no longer broadly coherent across the full network. This condition replicates the difficulties observed in recording persistent OB gamma in slices using single low-impedance glass LFP electrodes (Bathellier et al 2006, Lagier et al 2004). However, when field potentials were measured separately at individual electrode sites in the slice model, reflecting MEA recordings, strong oscillations were recorded, and regions of coherence similar to those found in slice recordings were observed. Reference electrodes were identified at three distinct locations and frequencies using the same methods described for slice recording analyses (Fig. 7*D*). At each frequency (44, 35, 37 Hz), a local coherence region was observed across several electrodes adjacent to the reference electrode for that frequency, whereas more distant electrodes were not coherent with the reference electrode. When longer-distance connections were restored, regenerating the *in vivo* condition, the entire network oscillated at a single dominant frequency of 43 Hz. Pairwise coherence patterns then were calculated based on the same three reference electrodes defined in the slice simulation; all three analyses demonstrated that the entire network was strongly and uniformly coherent, without regard to the physical distances between columns (Fig. 7*E*). Broad and stable coherence therefore can be generated in large-scale mixed networks in which the excitatory population exhibits intrinsic oscillations (Desmaisons et al 1999), but in which these local dynamics do not directly drive the network oscillation. This biophysical modeling result reflects the theoretical capacity of coupled-oscillator systems to synchronize extensive networks with heterogeneous coupling properties and relatively sparse small-world topologies (Kuramoto 1975, Li & Cleland 2017, Mallada & Tang 2013, Strogatz & Mirollo 1988, Watts 1999).

## Discussion

We here report a dynamical mechanism for robustly generating zero-phase coherence across a heterogeneously-activated population of principal neurons, instantiated in the neural circuitry of the OB. Using multielectrode array recordings from OB slices in which we induced persistent gamma-band oscillatory activity with neurochemical or optical stimulation, we showed that the OB network comprises multiple independent columnar oscillators that, while individually GABA(A) receptor-independent, are coupled together by GABA(A)-dependent synaptic interactions. We have examined the properties of this resonance-enhanced network gamma mechanism, pyramidal resonance interneuron network gamma (PRING), in related theoretical work (Li & Cleland 2017). Comparing intracellular recordings from the OB in slices and *in vivo*, while using spatiotemporally patterned optical stimulation, we showed that, in the intact OB, MCs exert effects on GCs across essentially the entire OB via their extensive lateral dendrites, whereas GC effects on MCs were local, as has been theoretically proposed (Cleland & Borthakur 2020, Imam & Cleland 2020, McIntyre & Cleland 2016, McTavish et al 2012). In contrast, in OB slices, MC effects on GCs also were spatially limited, owing to the transection of most long-distance lateral dendritic connections. We then constructed a biophysically realistic computational model of the EPL network in order to test whether the elimination of these longer connections would decouple a fully coherent OB network into multiple regional oscillators, and, conversely, whether a model of OB slice activity replicating the data of Fig. 4*C* would generate globally coherent gamma oscillations based solely on the addition of longer-distance connections between MCs and GCs. The simulations confirmed these hypotheses (Fig. 7), demonstrating that the observed pattern of regionally coupled oscillators in MEA slice recordings is a predictable outcome of slice preparation from an *in vivo* network of this type that exhibits global coherence (Ermentrout & Kopell 1991, Li & Cleland 2017, Rulkov & Bazhenov 2008).

We found that the blockade of GABA(A) receptors in the OB does not reduce the power of gamma oscillations (Fig. 5), a result superficially contrary to prior results (Bathellier et al 2006, Lagier et al 2004). We attribute this critical difference to the properties of the different electrodes used. Larger, lower-impedance LFP electrodes detect electrical potential changes over broader areas, spatially averaging across this larger “pickup radius” and thereby smoothing out highly local events (Viswam et al 2019). In studies using such electrodes, the loss of intercolumnar coherence that we observed owing to decoupling by GABA(A) receptor blockade would be detected as a loss of oscillatory power, and indeed was interpreted as such by these authors. In contrast, our spatially distributed, higher-impedance MEA electrodes each recorded from much more localized regions of tissue (i.e., smaller pickup radii), as evidenced by the fact that LFP recordings from neighboring electrodes could be decorrelated by pharmacological treatments (revealing that neighboring electrodes are not predominantly recording electrical fields generated from a common source location). This is corroborated by the observation that oscillatory power did not decrease after GABA(A) receptor blockade, implying that the total power of the (spatially averaged) field recorded by a given electrode was not drawn appreciably from regions of tissue outside of the rough scale of a “column”.

The GABA(A) receptor-independent local oscillations that we observed (Fig. 5) are likely to reflect the intrinsic subthreshold oscillations (STOs) of sibling MCs within an MOB column (Brea et al 2009, Desmaisons et al 1999), perhaps rendered locally coherent via intraglomerular gap junction coupling (Christie et al 2005, Friedman & Strowbridge 2003, Migliore et al 2005, Pouille et al 2017). In the absence of the intercolumnar connectivity mediated by GABA(A)ergic synapses, these intracolumnar oscillators drift in phase with respect to one another, becoming incoherent as observed herein (Fig. 5*C-G*). In the presence of this intercolumnar connectivity, however, MC STOs still contribute to the robust, zero-phase global coherence that the OB appears to exhibit (Li & Cleland 2017) and which is critical to contemporary theories of spike timing-based computation (Imam & Cleland 2020). It is established that coupled-oscillator dynamics are capable of globally synchronizing spatially extensive networks (Winfree 2001), and that this global synchronization capacity can extend to networks with sparser, small-world coupling topologies, different coupling strengths and functions, and heterogeneous coupling delays (Burton et al 2012, Li & Cleland 2017, Watts 1999, Zhou et al 2013). Briefly, in the OB, shunting inhibitory synaptic inputs reset the phase of MC STOs (Desmaisons et al 1999, Galan et al 2005, Galán et al 2006, Rubin & Cleland 2006). MCs receiving synchronous GABA(A)-ergic inhibition from GCs consequently will be reset to a common STO phase. The spikes generated by excitatory (afferent) inputs to MCs, in contrast, are temporally constrained by STO phase (Desmaisons et al 1999, Rubin & Cleland 2006). Differences in sensory excitation levels among MCs consequently will shape the timing of MC spike initiation and propagation, with stronger afferent inputs likely to yield spike phase advances within the broader phase constraints imposed by STO dynamics (*phase coding*) (Li & Cleland 2013, McIntyre & Cleland 2016). In this framework, ongoing network gamma oscillations depend on cycle-by-cycle STO resets by shunting inhibition, with newly-reset STOs constraining MC action potentials to a common phase window; these quasi-synchronized MC spikes then in turn evoke synchronous feedback inhibition from GCs, which again reset the STO phases of MCs within the activated ensemble (Li & Cleland 2017). The faster network gamma frequency exhibited when GABA(A)-ergic synapses are intact (Fig. 5*B*) arises from this interplay (Schoppa 2006), with the frequency depending primarily on the speed of inhibitory synaptic feedback, which governs the rate at which MC STOs are repeatedly reset. When this inhibitory feedback loop is blocked (Fig. 5*B*), the local oscillations are reduced to the slower intrinsic frequencies of MC STOs, which are voltage-dependent but generally in the beta band (Desmaisons et al 1999). The dynamics of these interactions have been explored in related theoretical work (Li & Cleland 2017).

A corollary of this coupled-oscillator framework is that gamma dynamics are intrinsic properties of the OB network architecture, such that any input providing overall excitation to principal neurons is likely to generate gamma oscillations with common characteristics. Afferent excitation, here mimicked by the optogenetic stimulation of primary receptor neuron axonal arbors, synaptically activates external tufted (ET) cells as well as other (inhibitory) glomerular-layer interneurons, and thereby excites mitral and tufted principal neurons (Gire et al 2012). Nicotinic cholinergic neuromodulation directly excites ET cells and mitral/tufted principal neurons (D’Souza et al 2013, Liu et al 2015, Rothermel et al 2014), whereas muscarinic effects potentiate the recurrent effects of glomerular-layer inhibitory interneurons (Liu et al 2015) and increase excitability in GCs (Smith et al 2015). Overall, the excitation of OB circuits via cholinergic modulation resembles that caused by afferent activity, particularly when only direct cholinergic effects upon OB circuitry are considered (Rothermel et al 2014). In contrast, metabotropic glutamate receptor activation is well known to evoke gamma oscillations in hippocampal circuits (Martin 2001, Whittington et al 1995), but is less well studied in the OB. In the OB, the activation of group I mGluRs excites ET cells (Dong & Ennis 2014, Dong et al 2009), generating afferent-like excitatory drive, while also inhibiting these cells via the activation of inhibitory interneurons (Dong et al 2007). Group II mGluR agonists both reduce activity in ET and mitral cells (Dong & Ennis 2017) and disinhibit these cells’ responses to strong afferent inputs (Dong & Ennis 2017, Zak et al 2015), on balance enhancing the contrast between strong and weak inputs. Importantly, under natural conditions, the pattern of activation of these mGluR receptors would be closely tied to locally-governed glutamate release and the concomitant effects of AMPA and NMDA receptor excitation. Our results suggest that, while mGluR activation does evoke OB gamma oscillations via its predominantly excitatory effects on the circuit, the overall combination of effects is less well coordinated with the functional circuit than those resulting from cholinergic neuromodulation or afferent activity, leading to a greater variability in response properties across slices (e.g., Fig. 2B).

As our understanding of the biophysical mechanisms underlying computations in neural systems becomes more sophisticated, transiently coherent activity within and between brain structures is increasingly implicated in normal brain function (Fries 2009, Fries 2015, Fries et al 2008, Kay 2014, Vinck et al 2010, Womelsdorf et al 2012). To understand the mechanisms by which coherence patterns are managed and shifted according to environmental, state, or task parameters (Frederick et al 2016, Fries 2015), and how they reflect underlying spike timing-based computations and the gating of interareal communication, will require the careful synthesis of diverse experimental investigations and theoretical analyses that incorporate the biophysical properties and limitations of neural circuits.

## Acknowledgments

The authors thank Dr. Jenny Davie for establishing patterned optogenetic illumination *in vitro* and support of the *in vitro* experiments, Drs. Venkatesh Murthy and Tom Bozza for providing genetically modified mice, and the international NSF/BMBF Collaborative Research in Computational Neuroscience program, the U.S. NIH/NIDCD, and the Max-Planck-Society for supporting this work.

---

**Figure S1.**
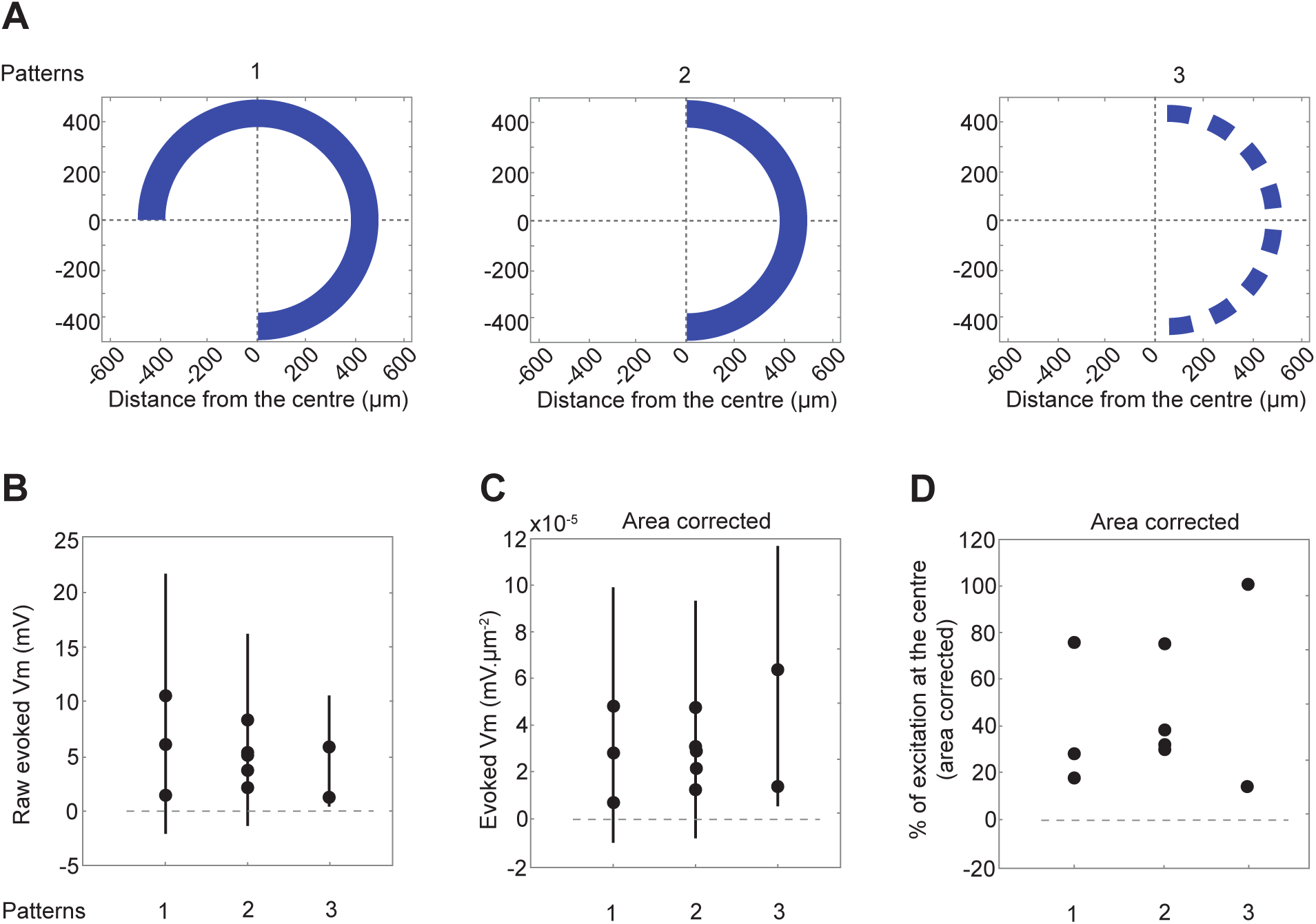
Measuring the effect of total luminosity. In the in vivo experiments of Fig. 6D-F, the concentric ring stimuli illuminated larger total areas and delivered higher total luminosity as the radius of the ring increased. The total luminosity could have an effect on the activation of granule cells, which (we show) receive excitation from illuminated areas distributed far across the OB. Accordingly, we assessed the impact of total luminosity decoupled from ring radius by comparing the effects of different light patterns on the depolarizations evoked in granule cells. In these experiments, three patterns of equal radius (400 µm) were used to stimulate the OB (patterns 1-3, in order of decreasing total area; panel A) while recording intracellularly from GCs (n = 3 cells). No light immediately anterior to the recording site was illuminated to avoid stimulating the passing olfactory nerve. The total luminosity was the highest for pattern 1, and lowest for pattern 3. The average change in the membrane potential relative to the baseline period is shown uncorrected (panel B), normalized by the total illuminated area (panel C), and further normalized by the depolarization following the illumination of the spot that maximally excited the recorded neuron (panel D). Uncorrected GC response amplitudes appeared to be affected by the total illuminated area (slope ∼10 µV/µm2), consistent with the finding that individual GCs receive and accumulate lateral excitatory input drawn from broad regions of the OB. This also may contribute to the slight (albeit not significant) tendency for the evoked responses of GCs to increase in response to illumination at greater distances (Fig. 6F). Vertical lines in panels B-C denote standard deviations.

